# Biologically Guided Physics-Informed Neural Network for Personalized PSA Forecasting in Intermittent Prostate Cancer Therapy

**DOI:** 10.64898/2026.11.19.767900

**Authors:** Shreya Yashodhar, Kevin Leder

## Abstract

Intermittent Androgen Deprivation Therapy (IADT) offers a promising management strategy for prostate cancer by alternating treatment periods with the aim of delaying castration resistance and mitigating side effects. An evolutionary rationale for treatment interruptions is to preserve treatment-sensitive cells that may competitively suppress resistant populations. However, designing optimal patient-specific treatment schedules remains challenging due to the complex dynamics of tumor evolution and the computational intensity of traditional mechanistic simulation models.

Standard mathematical models have prespecified structures that may inadequately capture patient-specific tumor evolution, limiting Prostate Specific Antigen (PSA) forecasting accuracy and adaptation to new patient data. These limitations may reduce the effectiveness of treatment policies learned using these models. To address them, we present a domain-adapted machine learning framework that integrates biological constraints into the deep learning architecture and embeds historical biomarkers to act as a robust, computationally efficient neural surrogate for mechanistic simulations.

This framework enhances generalization across sparse clinical profiles by leveraging deep neural representations for patient-specific and cross-patient parameter learning, and potentially enables the identi-fication of personalized dosing schedules tailored to individual tumor trajectories. Preliminary evaluations confirm that it reliably captures treatment-associated PSA dynamics while overcoming the computational bottlenecks of numerical solvers. Ultimately, this work provides a scalable foundation for optimizing adaptive therapy protocols to extend time to disease progression. We present the architectural formulation, forecasting evaluation against mechanistic baselines, and training and optimization procedures of the deep neural networks.

## 1 Introduction

### 1.1 Prostate cancer dynamics and androgen deprivation

Prostate cancer is the second most commonly diagnosed cancer among men worldwide and a major cause of cancer-related mortality [9]. Tumor growth depends on interactions among cancer cells, their microenvironment, and treatment. Several evolutionary models approximate prostate tumors as two competing populations: treatment-sensitive and treatment-resistant cells [3, 6, 17]. These populations compete for shared resources and space, and their abundance, growth rates, and treatment responses influence disease progression. This two-population representation is a modeling simplification of the broader heterogeneity present in prostate cancer [3, 6].

Androgen-targeted treatment primarily suppresses cells that depend on androgen signaling for growth and survival (sensitive population). Although reducing the sensitive population can decrease total tumor burden, it may also relieve competitive pressure on resistant cells, allowing them to expand. This process is termed competitive release. Evolutionary adaptive therapy therefore aims to preserve sufficient sensitive cells to suppress resistant growth while maintaining clinically acceptable tumor burden. Its effectiveness depends on the strength of competition and the biological characteristics of the tumor [3, 12].

Resistance can arise through genetic alterations and non-genetic changes, including altered gene expression and phenotypic switching. Some non-genetic resistant states may be reversible, whereas others can facilitate persistent resistance [11, 18]. Intermittent treatment changes the selective environment over time, potentially allowing sensitive populations to recover during treatment holidays. However, the rationale for adaptive therapy does not require all resistance to be reversible: competitive suppression of pre-existing resistant cells is itself a central mechanism [3, 12].

### 1.2 Androgen signaling and treatment mechanisms

Testosterone and related hormones, called androgens, support the growth of many prostate cancer cells. Most testosterone is produced by the testes, while the adrenal glands produce other androgens that can be converted into testosterone. Some prostate tumors can also produce androgens. These hormones activate androgen receptors -proteins in cells that trigger signals for growth and survival [10].

Cyproterone acetate (CPA) and Leuprolide acetate (LEU) were the two primary androgen-blocking therapies administered in the Bruchovsky cohort and are therefore the treatments considered in this study. CPA blocks androgen receptors and can also reduce testosterone production. LEU causes a brief initial rise in testosterone, followed by sustained suppression of its production by the testes. Both these in-turn suppress hormone-sensitive cancer cells. However, some cancer cells become resistant by increasing their androgen receptors or changing how those receptors work, allowing growth despite low testosterone levels [10].

Following treatment withdrawal, testosterone may recover over time, potentially permitting sensitive cells to regrow and restore competitive pressure on resistant cells [1, 10]. Adaptive therapy aims to preserve these cells so that they compete with resistant cells for resources and slow their growth [3, 12].

### 1.3 PSA measurements and adaptive therapy

Prostate-specific antigen (PSA) is produced by both normal and malignant prostate cells and is used to monitor treatment response and disease progression. Serum PSA is an indirect biomarker rather than a direct measurement of cancer cell number, growth rate, or the sensitive-to-resistant population ratio. PSA production is also influenced by androgen receptor signaling [13, 16].

Intermittent androgen deprivation therapy (IADT) introduces treatment holidays, with treatment withdrawal and resumption governed by PSA measurements and protocol-specific criteria. Earlier protocols, including the Bruchovsky trial, combined prescribed treatment periods with predefined PSA thresholds [1]. Although these rules respond to PSA changes, their thresholds are not necessarily optimized for individual patients’ tumor growth rates, treatment responses, or resistance dynamics.

### 1.4 Personalization using mechanistic models of tumor evolution

Mechanistic models provide a framework for studying how treatment alters competition between sensitive and resistant tumor populations. Ordinary differential equation models such as the Lotka–Volterra (LV) tumor-cell competition models can represent proliferation, cell death, resource competition, and treatment response, allowing researchers to compare treatment schedules that have not been administered clinically [3, 6].

Zhang et al. investigated an adaptive abiraterone protocol informed by mechanistic simulations based on LV models. In a pilot clinical trial involving 17 patients, treatment was suspended after a 50% reduction in PSA from baseline and resumed when PSA returned to baseline [2, 8]. 4 patients remained without radiographic progression at the reported follow-up. Subsequently, longitudinal clinical data were used to update a two-population model of treatment-sensitive and resistant cells. Retrospective simulations suggested that more prompt treatment withdrawal upon reaching the PSA threshold could better preserve sensitive cells, suppress resistant populations, and prolong tumor control [8].

Strobl et al. developed a two-population LV model and fitted patient-specific parameters to longitudinal PSA measurements from the Bruchovsky cohort [1, 3]. Their analysis examined the interaction between resistance costs, cellular turnover, initial tumor density, and the initial resistant fraction. Estimation of parameters such as proliferation rate, death rate and cell density minimized the root mean squared difference between normalized PSA and normalized predicted tumor density. Consequently, this model was informed by actual patient data. However, reproducing observed trajectories does not itself establish accurate forecasting beyond the fitting period. Further, evaluating these models on held-out observations interspersed within the fitted trajectory primarily assesses interpolation, limiting the evidence for their ability to forecast future disease dynamics beyond the observed period.

### 1.5 Reinforcement learning for individualized treatment

Reinforcement learning extends this framework by using a simulator as an environment in which an agent learns treatment decisions [7]. Engelhardt demonstrated this principle using a stochastic simulator of bacterial growth and resistance, parameterized with illustrative biological values rather than patient-specific data [4]. Such studies establish computational feasibility, while their treatment outcomes remain conditional on the simulated dynamics.

Gallagher et al. used the simulated trajectories from the patient-calibrated LV model from Strobl et al. to investigate personalized adaptive treatment [5]. Their framework learned treatment decisions from total tumor burden and explored both individual and randomized group training. They also investigated personalization from an initial treatment cycle. Importantly, the authors acknowledged that some fitted virtual patients failed to reproduce clinically observed progression, and that first-cycle fits could inadequately capture later dynamics. These findings underscore the distinction between policy performance within a simulator and the simulator’s ability to represent future patient trajectories.

In a related approach, Lu et al. developed a time-varied, mixed-effect generalized LV model incorporating patient-specific tumor dynamics and the effects of cyproterone acetate and leuprolide acetate [6]. Their I^2^ADT framework used this simulator to train individualized treatment policies. Simulator validation randomly withheld 20% of observations within each treatment cycle and reported an overall PSA (R^2^) of 0.84. This evaluates prediction within the observed treatment history rather than a chronological forecast beyond a cutoff.

### 1.6 Motivation for the present method

These studies motivate evaluating the forecasting ability of patient simulators separately from the treatment policies trained on them. Our study compares mechanistic ODE models with physics-informed neural networks [14] under matched chronological evaluation: models use an initial portion of each trajectory and forecast subsequent PSA without receiving held-out PSA observations.

We investigate whether neural models can represent patient dynamics inadequately captured by the selected ODE formulation. This includes whether population pretraining improves adaptation when individual histories are limited, whether patient embeddings capture useful differences in treatment response and whether retrospective fine-tuning using an initial chronological portion of each patient’s observations can improve patient-specific forecasts. These questions are investigated through comparisons of individual and population PINN configurations with the retrained ODE baseline. Improved PSA forecasting would establish a robust forecasting foundation for subsequent treatment-policy optimization.

## 2 Methods

### 2.1 Experiment design

We evaluated physics-informed neural networks (PINNs) trained on population and patient-specific histories to forecast longitudinal prostate-specific antigen (PSA) dynamics. Multiple configurations were developed to assess their suitability as neural surrogates for the time-varied mixed-effect generalized Lotka–Volterra (tM-GLV) model. PSA and testosterone were the available longitudinal biomarkers; PSA served as the forecasting target, while testosterone was used to characterize patient cohorts. The framework combines observed patient histories with mechanistic constraints on intratumoral competition to learn latent patient characteristics and model individualized dynamics, providing a foundation for a patient-specific state-transition simulator for intermittent androgen deprivation therapy optimization.

The experiments examined individual-patient learning, population learning, roll-forward forecasting, and adaptation to patients excluded from population pretraining. A chronologically retrained time-varied mixed-effect generalized Lotka–Volterra (tM-GLV) model served as the mechanistic comparator. One-step prediction and recursive forecasting were evaluated separately because they use different amounts of observed patient information.

The study focused on the predictive component of a potential reinforcement-learning environment. Treatment-policy learning and optimization were not evaluated in these experiments. Forecasts were conditioned on recorded treatment histories, allowing the models to be compared against subsequently observed PSA under the treatment schedules actually administered. Extending it to PSA-guided treatment optimization will require sequentially applying treatment decisions based on predicted PSA and forecasting the resulting PSA trajectory under those decisions.

### 2.2 Clinical data and preprocessing

Longitudinal data were obtained from the Bruchovsky intermittent androgen deprivation therapy cohort [1]. Dataset characteristics and preprocessing procedures are summarized in Table 1.

**Table 1.** Clinical dataset characteristics and preprocessing procedures.

| Item | Description |
|---|---|
| Available records | Patient identifiers, observation times, PSA, CPA and LEU dosages, testosterone, treatment-cycle information, and treatment status. |
| Study cohort | The dataset contained 72 patients. Four patients with fewer than 10 visits following trajectory truncation at medication switching were excluded, leaving 68 patients. |
| Chronological ordering | Observations were ordered chronologically within each patient. |
| Forecasting target | Observed PSA. Records without measured PSA were excluded. |
| *Cohort classification (cohort sizes are reported separately for each analysis)* |  |
| Train & validation | After chronological train-validation splitting, eligible data points for each experiment required enough visits to form the multi-step prediction window (for roll-forward<sup>1</sup> training) and evaluate predictions on subsequent visits. Minimum visit thresholds ranged from 3 to 10 across experiments. |
| History window | The patient-embedding history window ranged from 6 to 8 visits across experiments, with padding used when fewer prior visits were available. |
| Descriptive variables | Testosterone and treatment-cycle summaries were used to characterize patient and cohort distributions. |
| *Neural network* |  |
| Visit-level predictors | Time, preceding PSA, and preceding CPA and LEU administration. Treatment inputs were binary indicators of whether the preceding dose was positive. |
| Input & target scaling | Time was scaled to represent years and in the roll-forward models<sup>1</sup>, preceding & target PSA were scaled by dividing by 40 before training. |
| Mechanistic treatment inputs | Treatment-response arrays were constructed over intervals between visits and retained for the mechanistic component. |

### 2.3 Mechanistic baseline and constraints

The mechanistic baseline used the time-varied mixed-effect generalized Lotka–Volterra (tM-GLV) model developed by Lu et al. [6]. A time-varying competition matrix allows interactions between the populations to change during disease evolution.

Each individual patient’s model represents two competing tumor populations *x*_1_ and *x*_2_, being the androgen-dependent and androgen-independent populations respectively, and *p* the PSA. Their dynamics depend on intrinsic growth rates, carrying capacities, competition for shared resources, and treatment-induced suppression. While initial values of *p* are available, initial values of *x*_1_ and *x*_2_, computed as fixed proportions of starting carrying capacities *K*_1_ and *K*_2_ respectively, are estimated using Lu’s assumptions [6]. Initial values of *x*_1_ and *x*_2_ are computed as

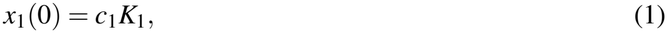

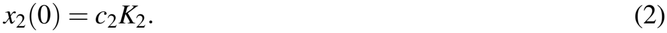

For estimating the carrying capacities, the mean volume used is 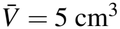, the mean pretreatment PSA is P = 22.1 *µ*g*/*L, and the cell size is *V*_cell_ = 5.236 *×* 10^*™*10^ *cm*^*3*^. Average pretreatment PSA is from the Bruchovsky trial, whereas mean volume was derived by Lu assuming that 50% of the average prostate volume decline in the Bruchovsky clinical trial [1] is attributable to cancer cells. Lu et al. assumed a cancer-cell radius of 5 *µ*m and computed the volume of a single cell as below [6]:

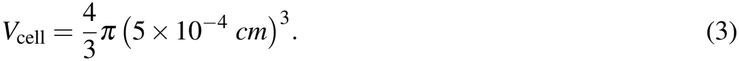

Subsequently, carrying capacities were computed as

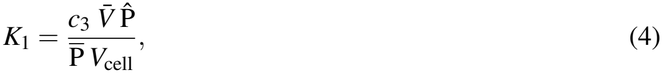

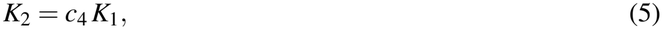

which, with the values above, become

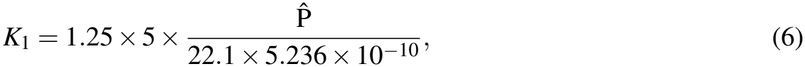

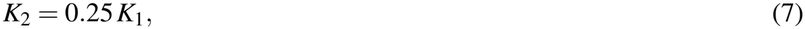

where *K*_1_ and *K*_2_ represent the carrying capacity of the androgen-dependent and androgen-independent cells respectively. 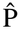 is the maximum empirical PSA level of the training data for each individual patient, so **K** = (*K*_1_, *K*_2_) is fixed and is not a learned parameter. Time in the competition term is in years, *t*_*y*_ = *t*_days_*/*(28 *×* 12).

CPA and LEU administration enter through drug-response functions representing their effects over time. The ODE sees a continuous daily drug-response series, built the same way as in Lu et al. [6] CPA on a visit is held at dose/200 until the next visit. A first LEU injection of 7.5, 22.5, 30, or 45 mg is mapped to a piecewise ramp over 4, 12, 16, or 24 weeks: a linear flare over days 0–6, a rise through day 20, then a plateau at the normalized depot^1^ level. A repeat injection within seven days of the previous depot’s end holds the response at that plateau. The response is divided by the nominal dose so that a full depot is 1.

A coupled PSA equation links the latent tumor populations to measured serum PSA through production and clearance terms, allowing patient-specific parameters to be estimated from longitudinal PSA and treatment records. The state equations themselves are in days. With clearance *φ* = 0.25,

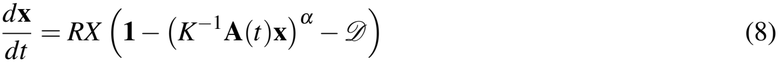

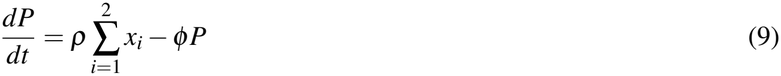

where R=[*r*_1_, *r*_2_] represents a vector of growth rates of the two phenotypes, *α* is the constant, hyperpa-rameter of competition-induced decay, *D* is the drug term and *ρ* is the PSA secretion rate. The drug term is *D* = *β* **d**(*t*), where the drug response vector is **d**(*t*) = (*d*_CPA_, *d*_LEU_) and *β* is (*β*_1_, *β*_2_)^*⊤*^. The competition matrix *A* has ones on the diagonal and off-diagonal entry 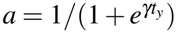. The initial state is *x*_1_(0) = 0.8 *K*_1_, *x*_2_(0) = 10^*™*5^ *K*_2_, based on the assumption that initially responsive cells constituted the majority of the tumor microenvironment, and *p*(0) equal to the observed baseline PSA. The coupled equations link latent tumor-population dynamics to observed PSA through modeled production and clearance [6].

The fitted ODE uses Lu’s seven-parameter vector Θ = (*r*_1_, *r*_2_, *β*_1_, *β*_2_, *α, γ, ρ*). The ODE model estimated the parameters through a forward method of numerical integration using a xitorch solver, and backpropagation through the loss between the predicted and observed PSA to update the parameters (Figure 1).

**Figure 1.**
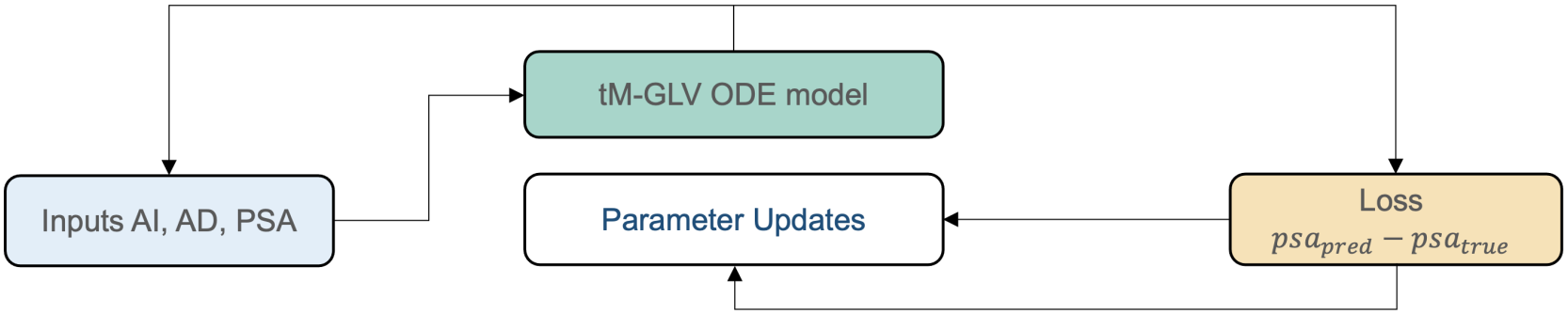
Training workflow of the mechanistic tM-GLV baseline. Patient inputs (androgen-dependent and androgen-independent populations, AD and AI, and PSA) are propagated by forward ODE integration; the loss between predicted and observed PSA is backpropagated to update the ODE parameters, and the predicted PSA is fed back as the next input.

### 2.4 Mechanistic baseline retraining

The tM-GLV implementation associated with Lu et al. [6] was retrained separately for the ten comparison patients. The original random withholding of observations was replaced by a chronological division: the first 80% of observations were used for fitting and the final 20% for evaluation. Patient-specific parameters were estimated from the fitting interval, and subsequent PSA was predicted by continuing the model forward under the recorded treatment schedule.

Piecewise numerical integration was retained across treatment intervals to preserve changes in treatment exposure. The final state of one interval provided the initial state for the next, maintaining continuity of the modelled trajectory. Held-out PSA measurements were used to score predictions rather than to reset the forecast state.

### 2.5 Neural representation

The PINN architectures combined time, normalized preceding PSA, treatment information, and, in embedding-enabled configurations, a learned representation of patient history. The treatment information includes binary indicators of whether the previous visit’s LEU and CPA doses were positive. Historical observations were organized into fixed-length windows so that the patient representation could summarize recent PSA and treatment patterns.

For visit *i*, the network receives scaled time, preceding treatment indicators, preceding PSA, and a patient representation:

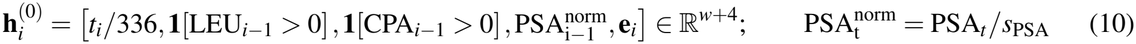

where *t*_*i*_ is elapsed time in days, *s*_PSA_ is the fixed PSA scaling constant, **1**[*·*] denotes the indicator function, and **e**_*i*_ ∈ R^*w*^ is the learned patient representation with embedding dimension *w*, ranging from 6 to 8 across experiments.

The input is propagated through three hidden layers with hyperbolic tangent activation:

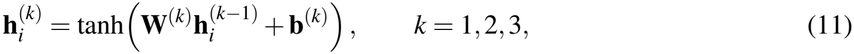

where **W**^(*k*)^ and **b**^(*k*)^ are the trainable weight matrix and bias vector of layer *k*. A linear output layer produces three unconstrained outputs:

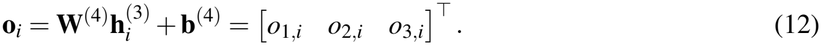

Exponential transformations yield positive latent tumor-population states and normalized PSA predictions:

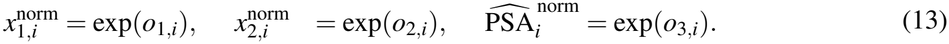

Here, 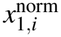 and 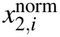 represent the modeled treatment-sensitive and treatment-resistant populations, respectively. Predicted *x*_1_, *x*_2_ and PSA are recovered on their original measurement scales as

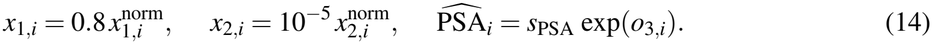

In the population models described in 2.7, patient representations provided a mechanism for conditioning these shared parameters on differences in individual trajectories. Patient representations were encoded using a Gated Recurrent Unit architecture [15], using a history window of 6 previous visits. To establish baseline feasibility, we use the GRU history encoder to derive a patient embedding from observed PSA values and condition the PINN on that same embedding throughout the entire patient trajectory.

When a history vector *z* is used, the biological parameters are Θ(*z*) = *µ* + Σ*z*, representing a mapping from patient states. Σ is initialized at zero, and *µ* is initialized so that training starts at the parameter means reported by Lu et al. [6]. Θ = (*R, D, γ, ρ, α*) with *R* = (*r*_1_, *r*_2_) and *β* = *β*_1_, *β*_2_

The physics component of the model used a time-modified Generalized Lotka–Volterra description of competing treatment-sensitive and treatment-resistant tumor populations, with the same initialisations as the mechanistic model. Growth, interpopulation competition, carrying capacity, and treatment effects constrained the modelled dynamics. PSA was the observed biomarker, while tumor-population states remained latent quantities inferred under the model assumptions, similar to the mechanistic model.

Parameter constraints on the values of (*r*_1_, *r*_2_, *β*_1_, *γ, ρ*) are borrowed from Lu et al. [6], and those on (*β*_2_, *α*) are biologically feasible constraints. Sigmoid and softplus functions are used to constrain the values between the identified limits.

The computation of *φ* , *K*_1_, *K*_2_ and *a* are a similar to the mechanistic model.

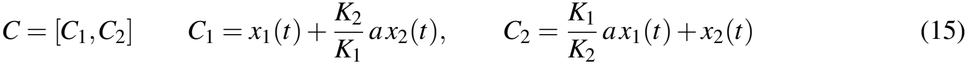

where *x* _*j*_ is as defined in eq. (14).

The biological constraints from the tM-GLV model, when applied on the output of the neural network are as:

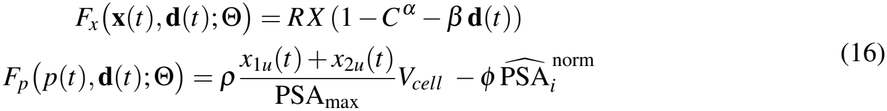

Where

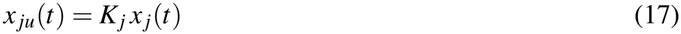

**d**(*t*) denotes treatment information, **x**(*t*) = [*x*_1_(*t*), *x*_2_(*t*)] is the vector of the population states, Θ is a representative term for mechanistic parameters, 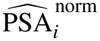 is from eq (13) and *x* _*ju*_(*t*) are the denormalized population values.

Physics residuals were evaluated at collocation points as the deviation of the automatic differentiation of neural predictions with respect to time 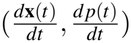 from the ODE differential in eq (16) [14]. For modelled states **x**, *p* and mechanistic dynamics *F*, a residual had the form

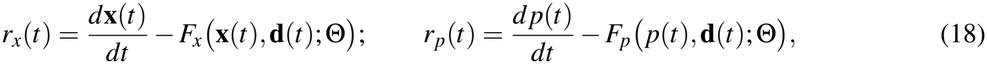

The physics loss guides neural predictions consistent with the biological formulation while retaining flexibility to learn patient-specific dynamics (Figure 2).

**Figure 2.**
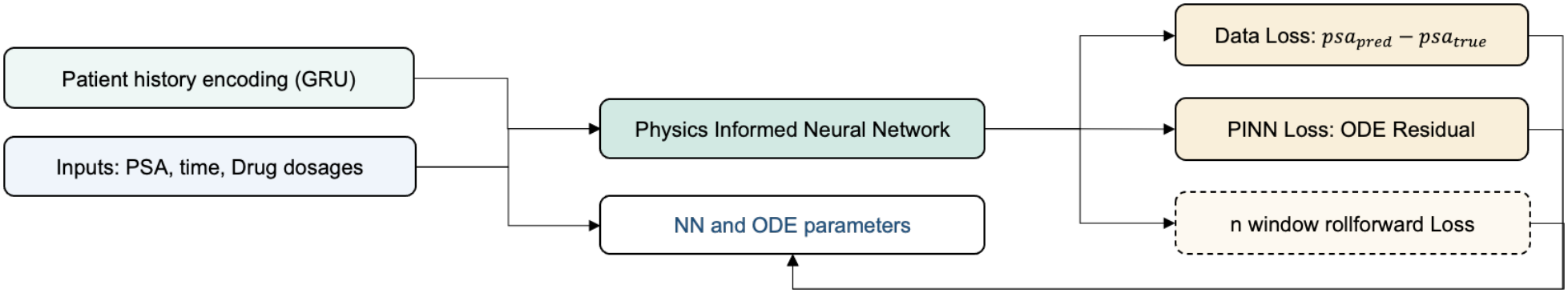
Training workflow of the physics-informed neural network (PINN). A GRU encodes patient history into an embedding that is supplied to the network together with the PSA, time and drug dosages. The data loss and the PINN (ODE-residual) loss are backpropagated to update the network and ODE parameters. The *n*-window roll-forward loss is backpropagated only in the roll-forward model.

### 2.6 PINN Training objectives and forecasting comparison

The combined training objective includes three components – the supervised data loss measured mean squared error between predicted and observed scaled PSA, a weighted physics-residual term [14] and the rollout loss.

The total loss is

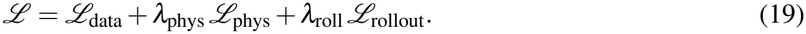

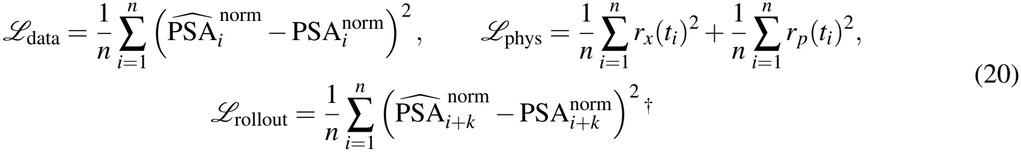

For the roll-forward objective, 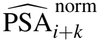 is the H-step prediction where output of the k-1 th step is the input of the kth step.

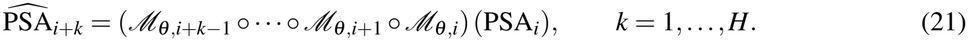

The physics weight was scheduled during training, with loss balancing used in the explored configurations to manage differences in the magnitudes of the two terms. Initial-condition warm-start training was explored to anchor the modelled state to the beginning of a patient trajectory and initial values of *x*_1_ and *x*_2_ detailed above. The network and mechanistic parameters were learned by minimizing the total loss function.

### 2.7 Model variants

Initial experiments were in the one-step setting, where each prediction used the preceding observed PSA and historical treatment information (Figure 2). Thus, observed PSA continued to be available as an input during one-step evaluation as detailed in Figure 3a. Under this input approach, single-patient approaches fitted a separate model to each patient, whereas the population approaches shared neural-network parameters across patients. This setting assessed prediction of the next observation and was distinguished from an uninterrupted forecast.

**Figure 3.**
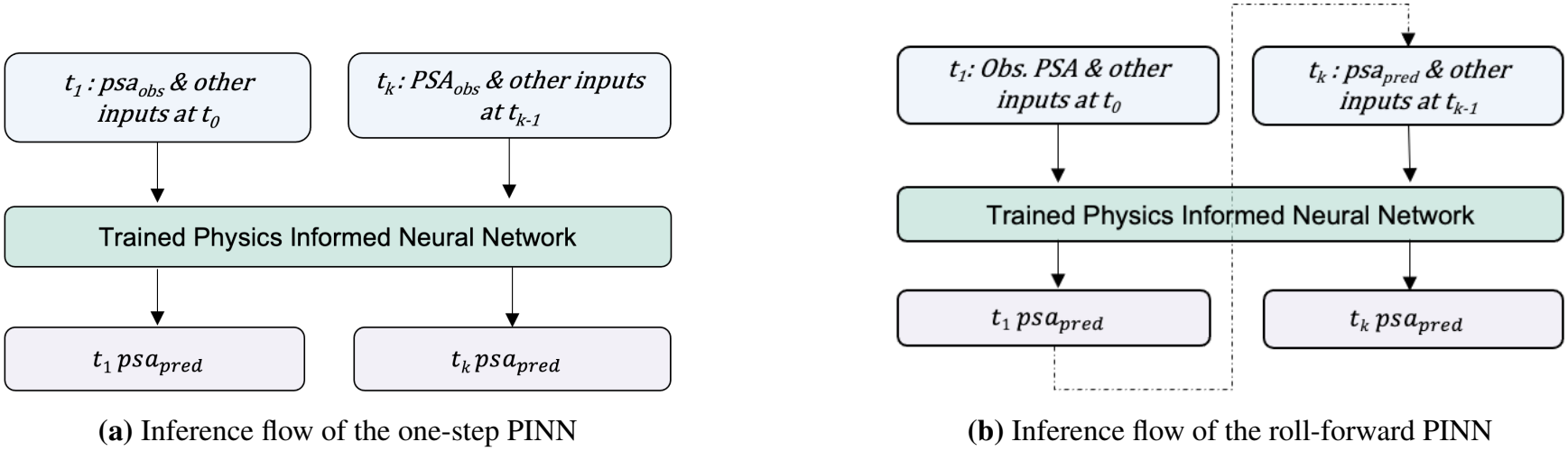
Inference flow of the physics-informed neural network (PINN). For the one-step PINN observed PSA at each timestep is fed as input while for the roll-forward PINN it is fed once at the first validation/test timestep and the prediction at timestep k is used as input for timestep k+1.

Subsequently,we performed roll-forward training, where predictions were propagated across successive visits, and errors were evaluated over multiple forecast steps (Figure 2). At evaluation, recursive forecasts were continued without supplying the held-out PSA measurements as demonstrated in Figure 3b. Recorded treatment information remained available. This setting assessed the accumulation of prediction error over time under the observed treatment schedule. Rollout loss used is the residual between the predicted PSA from continuous rolling predictions n steps from the current timestep and the observed PSA at that time (eq. 20).

Multiple neural configurations were explored, including patient embeddings, history-window length, network dimensions, scaling choices, and physics-loss weighting. Training used minibatches, learning-rate scheduling, and gradient clipping. Configuration comparisons were based on validation performance, with the selected configurations then evaluated over the corresponding reported intervals.

### 2.8 Patient splits and population adaptation

In the initial single-patient experiments, a separate model was trained for each patient using the first 80% of their observations and validated on the final 20%. Similarly, initial one-step population training used the 80% of the training patients’ visit information for training and the final 20% of their visit information for validation, with shared model parameters learned across these patients. The population one-step model was evaluated across the full trajectories of held-out test patients.

For the roll-forward experiments, 10 patients were withheld from population pretraining. We evaluated five variants. (i) The sequentially sampled model used the first 52 patients in the ordered pool for population training and the remaining 6 for validation, then was fine-tuned on the first 70% of each held-out patient’s observations. (ii) The statistically sampled model used treatment exposure, testosterone summaries, and treatment-cycle information to stratify patients into population training and validation groups, then was finetuned on the first 70% of each held-out patient’s observations. (iii) The Bruchovsky-sampled model formed population training and validation groups using the clinical response categories reported by Bruchovsky et al. [1] (responsive, resistant, and resistant with metastasis), then was fine-tuned on the first 70% of each held-out patient’s observations. (iv) A second statistically sampled variant used the same stratification strategy as (ii) and was fine-tuned on the first 50% of each held-out patient’s observations. (v) The population-only variant used the statistically sampled population model directly, without patient-specific fine-tuning.

Each sampling strategy for the roll-forward models assigned 52 of the 58 available patients to population training and 6 to validation, using their full trajectories for both. For patient-specific adaptation, variants (i)–(iii) used a chronological 70/10/20 split and variant (iv) used a 50/30/20 split. The first block was used for fine-tuning, the middle block for model selection, and the final block for test evaluation. Split boundaries were calculated using the full visit count, including the initial visit, and rounded to whole visits. In roll-forward forecasting, the model starts from an observed PSA value and recursively feeds each predicted PSA into the next step while using recorded treatment inputs throughout.

The population model without fine-tuning served as a transfer baseline. Its forecasts were initialized at the first visit and rolled forward recursively, with the initial visit excluded from scoring. We evaluated this baseline over the full rollout and extracted predictions for the final 20% block for comparison with the fine-tuned models. These final-block predictions came from the original rollout and therefore reflected the forecast history accumulated from the first visit.

We used patient-level scores and trajectory plots to compare the PINN forecasts with those of the retrained tM-GLV model. Both models were evaluated on the final 20% of their respective trajectories.

### 2.9 Performance assessment

Predictive performance was assessed on the original PSA scale using the coefficient of determination (*R*^2^) and root mean squared error (RMSE). Metrics were reported separately for training, validation, test, and overall trajectory intervals where available. The initial visit, when used for initialization, was excluded from the corresponding prediction scores. Patient-level distributions were summarized using means, medians, and the proportion of evaluable patients with *R*^2^ above 0.8. Pooled scores were computed by concatenating predictions and observations across patients for the relevant interval; these were distinguished from averages of patient-level scores. Scores with an undefined *R*^2^ were excluded from *R*^2^ summaries.

Representative trajectory plots were inspected alongside treatment histories to assess the timing and magnitude of predicted PSA changes. Patient and cohort summaries of PSA, drug exposure, testosterone, and cycle counts were used to explore differences in performance across sampling strategies and between early and later trajectory segments.

## 3 Results

### 3.1 Single patient, population one step prediction and population roll-forward

To assess whether physics-informed neural networks (PINNs) could capture temporal patterns in PSA dynamics, we first evaluated separate single-patient models and a shared population model using one-step prediction (Refer 2.7). In the population model, 56 patients were used for training and 12 patients were used as held-out test sets. Multiple configurations were evaluated and the results reflect architecture and configuration improvements.

The lower mean relative to the median reflects the influence of low-scoring patients. Population training improved aggregate one-step validation performance across all metrics, motivating subsequent evaluation of roll-forward prediction and patient-specific adaptation.

Next, we trained a population PINN with a roll-forward loss^†^ using the same 56 training patients. The slight reduction in performance of the roll-forward model is suggestive of the trade-off between immediate PSA forecasting and the lookahead accuracy. Its performance nevertheless exceeds that of the selected single-patient configuration, supporting the contribution of shared population learning. The increase in pooled validation *R*^2^ from 0.47 to 0.51 indicates improved aggregate predictive performance. Overall results are highlighted in Table 2 and representative patient trajectories for 2 individual patients are presented in Figure 4.

**Table 2.** Single-patient and population PINN performance (one-step prediction and roll-forward training).

| Statistics | One-step single | One-step<br>population | Roll-forward<br>population |
|---|---|---|---|
| Mean patient full-trajectory *R*<sup>2</sup> | 0.61 | **0.73** | 0.67 |
| Median patient full-trajectory *R*<sup>2</sup> | 0.77 | **0.81** | 0.76 |
| Median patient validation *R*<sup>2</sup> | 0.29 | **0.61** | 0.50 |
| Pooled validation *R*<sup>2</sup> | 0.12 | 0.47 | **0.51** |
| No. of patients with validation *R*<sup>2</sup> > 0.8 | 9 | **19** | 14 |
| Pct. of patients with validation *R*<sup>2</sup> > 0.8 | 16% | **34%** | 25% |

**Figure 4.**
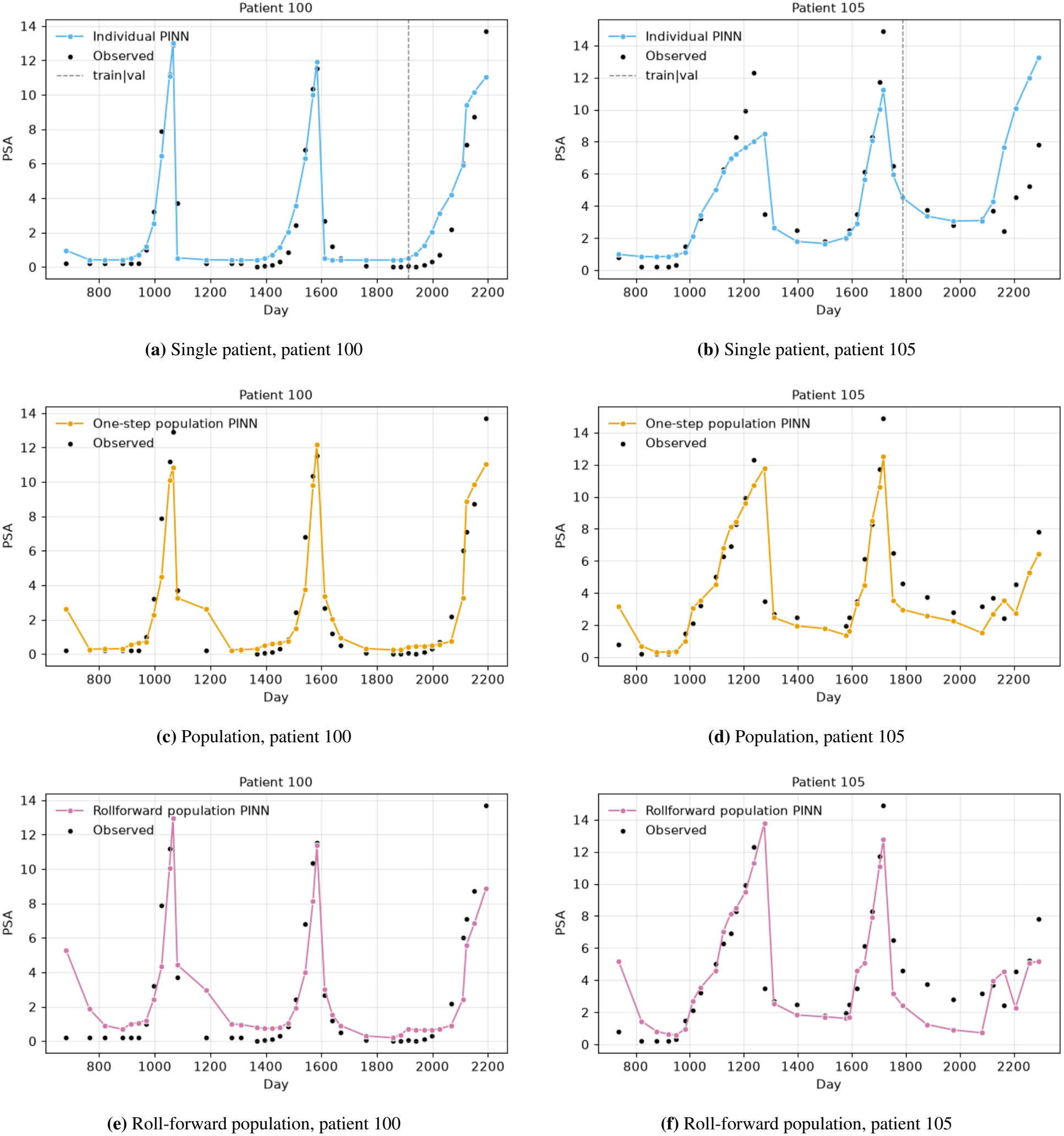
Representative trajectories for patients 100 and 105 illustrate the PSA patterns captured by the single-patient, population and roll-forward population predictions for patients 100 (left) and 105 (right). Validation *R*^2^ of patient 100 increased from 0.84 to 0.91 and reduced to 0.83 and similarly for patient 105 was -5.47, 0.74, 0.58 in the individual, population and roll-forward models respectively. Scores correspond to the evaluation intervals identified in each panel, as discussed in Section 2.8. The trajectories illustrate patient-specific differences in recursive forecasting performance, motivating adaptation using early observations from held-out patients.

### 3.2 Patient stratification and fine tuning

In the next step, we trained the roll-forward models using the various patient-split strategies as described in Section 2.7. The mean PSA, LEU and CPA for the population cohorts are presented in Table 3.

**Table 3.** Reported characteristics of population cohorts.

|  | Mean PSA (*µ*g/L) |  |  | Mean LEU (mg/d) |  |  | Mean CPA (mg/interval) |  |  |
|---|---|---|---|---|---|---|---|---|---|
| Cohort | Sequential | Statistical/<br>Population-only | Bruchovsky | Sequential | Statistical/<br>Population-only | Bruchovsky | Sequential | Statistical/<br>Population-only | Bruchovsky |
| Train | 4.48 | 4.39 | 4.35 | 1.95 | 1.99 | 1.92 | 47.25 | 46.43 | 45.67 |
| Validation | 3.54 | 4.35 | 4.66 | 2.36 | 2.00 | 2.60 | 41.74 | 48.89 | 55.42 |
| Test | 3.98 | 3.98 | 3.98 | 3.35 | 3.35 | 3.35 | 57.96 | 57.96 | 57.96 |

*Note:* visit-level averages for the various splits of training, validation and test sets. Drug dosage units are from the Bruchovsky dataset.

Table 4 summarizes overall trajectory fit and held-out test performance. The model trained using statistical sampling split of train/validation sets with the 70/10/20 split had the highest mean and median patient train + validation *R*^2^, *∼*0.73 and 0.80 respectively, whereas the statistical 50/30/20 configuration achieved the highest median test *R*^2^, both overall (0.78) and after excluding patients with relapse (0.80). If we exclude the patients with relapse (patients 11 and 12), the performance of both the statistical sample based training models and the population only model are fairly close.

**Table 4.**
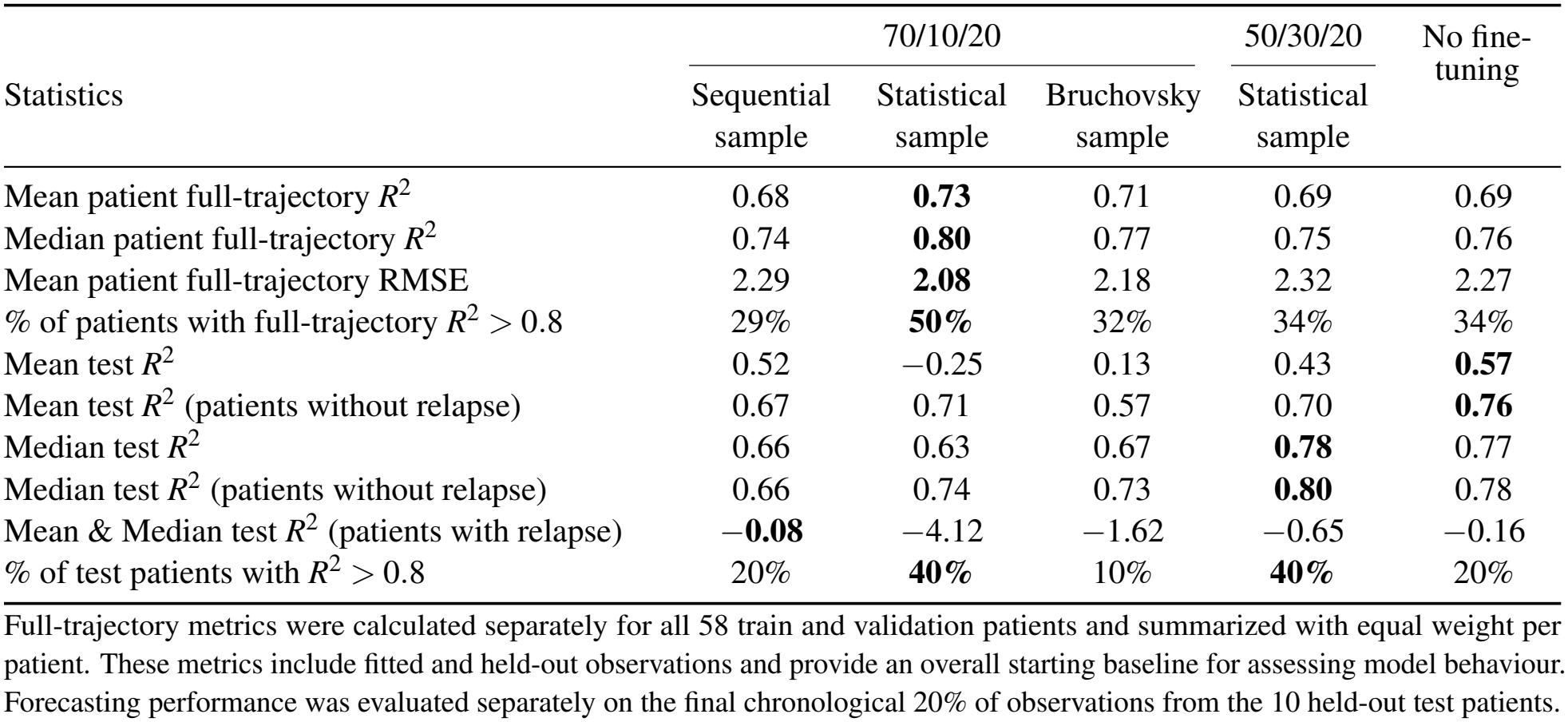
Overall and final test performance of population only and patient adaptation models.

The statistically sampled fine-tuned model achieved test *R*^2^ *>* 0.8 in 4 of 10 patients, compared with 2 for the sequential sample, 1 for the Bruchovsky sample and 2 without fine-tuning. However, its mean test *R*^2^ was *™*0.25 because severe failure in patient 12 outweighed improvements elsewhere. The frequency of strong predictions and the severity of failures therefore provide complementary information.

This indicates that 1) both statistical adaptation configurations produced 4 patients with test *R*^2^ *>* 0.8, while the population-only model achieved the highest mean test *R*^2^, motivating patient-specific selection between population forecasts and adaptation and 2) the variable adaptation outcomes in patients 11 and 12 indicate that fine-tuning requires particular attention when forecasting relapse-associated trajectories.

The complementary strengths of population forecasts and statistical fine-tuning support a patient-specific/ clustered adaptation strategy, rather than uniform fine-tuning across all patients.

### 3.3 Variation in patient specific performance

Table 5 compares patient specific performance using the final chronological test block for each approach. Population-only model scores were recomputed from the corresponding portion of the saved whole-trajectory rollout; the forecast was not restarted at the test cutoff.

**Table 5.** Patient-specific final test *R*^2^.

|  |  | Patient |  |  |  |  |  |  |  |  |  |
|---|---|---|---|---|---|---|---|---|---|---|---|
| Split | Sampling strategy | 3 | 12 | 87 | 91 | 15 | 37 | 58 | 11 | 77 | 100 |
| 70/10/20 | Sequential | 0.47 | 0.56 | 0.46 | 0.67 | 0.65 | 0.77 | 0.66 | −0.72 | 0.83 | 0.88 |
|  | Statistical | 0.50 | −8.21 | 0.51 | 0.60 | **0.88** | 0.83 | **0.90** | −**0.03** | **0.82** | 0.66 |
|  | Bruchovsky | 0.36 | −3.37 | −0.27 | 0.82 | 0.79 | 0.61 | 0.77 | 0.13 | 0.73 | 0.74 |
| 50/30/20 | Statistical | **0.76** | −0.07 | −0.09 | **0.82** | 0.80 | **0.91** | 0.81 | −1.23 | 0.66 | **0.91** |
| – | Population-only | **0.77** | **0.59** | **0.55** | **0.83** | 0.82 | 0.78 | 0.80 | −0.91 | 0.77 | 0.75 |

Statistical fine-tuning (70/10/20) improved test *R*^2^ relative to population-only in 5 of 10 patients; statistical fine-tuning (50/30/20) improved 3, sequential fine-tuning improved 3, and Bruchovsky sampling improved 1. These results demonstrate heterogeneous adaptation benefits. Identifying patient characteristics that determine which model performs best requires further investigation.

For patients 3, 12 and 87, mean PSA increased from 4.78 *µ*g/L, 3.31 *µ*g/L and 3.91 *µ*g/L during the fine-tuning training period to 6.69 *µ*g/L, 6.01 *µ*g/L and 7.07 *µ*g/L during testing. These distribution shifts over time could explain some aspects of model performance. Patient 12 illustrates the adaptation challenge most clearly: statistical and Bruchovsky fine-tuning produced test *R*^2^ values of *™*8.21 and *™*3.37, whereas sequential sampling and population-only achieved 0.56 and 0.59. In patient 91, mean CPA dosage declined from 67.65 to 45.45 between train and test periods.

Statistical fine-tuning with 70% of data used for fine-tuning produced strong test predictions for patients 15, 58 and 77, with *R*^2^ values between 0.82 and 0.90. The statistical PINN plots in Figure 5d, 5f & 5g show recurring PSA rises and declines, while remaining errors involve peak magnitude, decline timing and PSA levels between peaks. Together with the tM-GLV comparison, these results support our hypothesis that the biologically guided neural models can capture recurring PSA dynamics more robustly than the selected mechanistic formulation.

**Figure 5.**
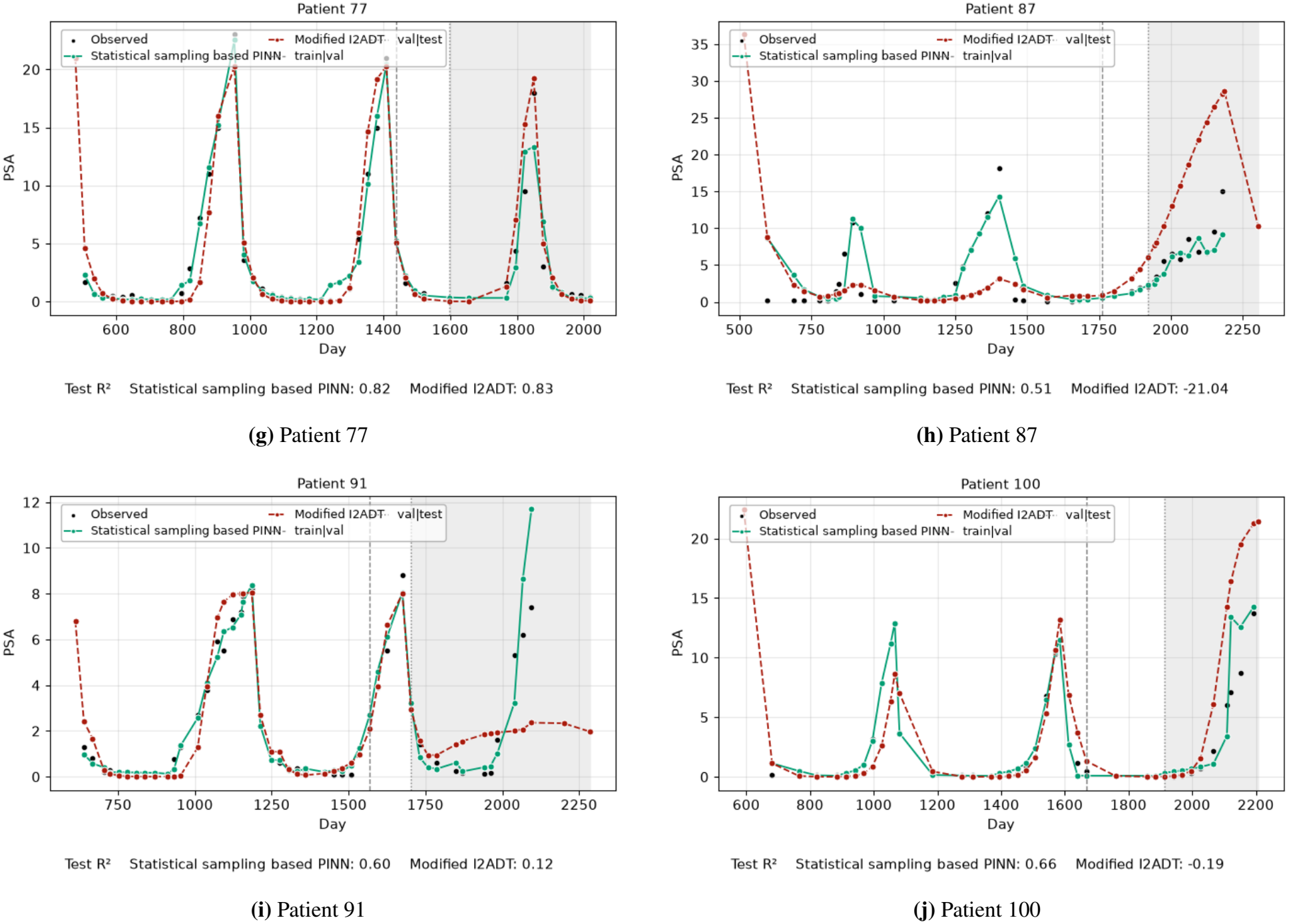
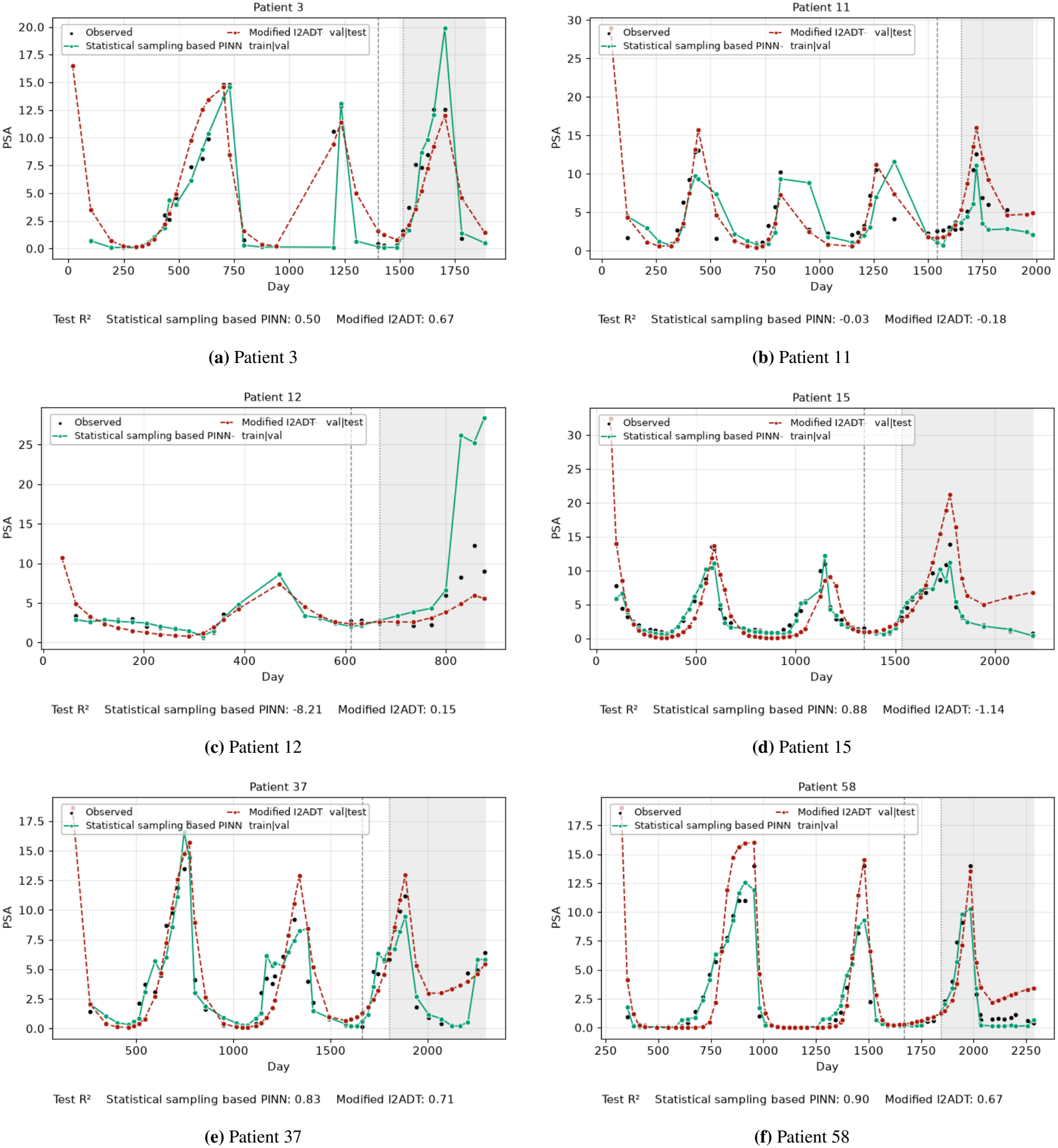
Patient-specific PSA trajectories for the ten held-out test patients.

Statistical fine-tuning with 50% of data used for fine-tuning performed best for patients 37 and 100, with test *R*^2^ of 0.91. Test-period mean PSA was 4.48 *µ*g/L for patient 37 and 4.31 *µ*g/L for patient 100, close to the population-training mean of 4.48 *µ*g/L. Both patients’ longer validation windows included a full PSA peak, providing additional trajectory information for model selection, possibly explaining the training success of this method for these patients.

Patient 11 remained difficult under every strategy, although some adapted models improved on the population baseline. Its plots show mismatches in decline timing and later PSA levels. Its short, nearly flat validation segment also produced extremely negative validation *R*^2^ values. Evaluation of models using longer validation windows to account for a later rise or treatment transition remains a challenge in the current dataset due to limited data availability.

These results motivate further exploration of optimal model choice at the intersection of the population only model and the % of chronological data used for training and validation in the adaptation phase, using patient-level cross-validation.

### 3.4 Comparison with the retrained tM-GLV model

Lu et al. calibrated their tM-GLV model by randomly withholding 20% of observations within each treatment cycle for validation [6]. This assesses prediction of missing observations within an observed trajectory. Their proposed personalized dosing policies were evaluated through model-based simulations informed by clinical data. To assess chronological forecasting of a comparative baseline, we retrained the tM-GLV baseline using the first 80% of each patient’s observations and reserved the final 20% for evaluation.

We compared the statistically sampled PINN with this retrained tM-GLV baseline. The PINN used 70% of observations for adaptation, 10% for validation, whereas tM-GLV used 80% for fitting; both were tested on the final 20%.

Figure 5 illustrates the supplied predictions.

The statistically stratified PINN achieved higher test *R*^2^ than the retrained tM-GLV baseline in 7 of 10 patients, closely matched performance in 1 patient and achieved lower scores in 2 patients (Refer Table 6). 4 patients achieved *R*^2^ *>* 0.8, compared with 1 for tM-GLV, demonstrating the framework’s potential to improve patient-specific PSA forecasting. Although adaptation was unsuccessful for patient 12 and both approaches struggled with patient 11, the overall results support biologically guided neural networks as a promising alternative to mechanistic models for capturing treatment-associated PSA dynamics. Further, these models train in approximately 20–30 minutes (including fine-tuning) for the population, in comparison with the *∼*22 hour timelines of the mechanistic models for each individual patient, supporting more frequent patient-specific model updates using visit-level data. These findings motivate evaluation across the full cohort and provide a foundation for extending the framework to adaptive treatment optimization.

**Table 6.** Held-out test *R*^2^ per patient: statistically sampled PINN (70/10/20) versus the chronologically retrained tM-GLV baseline.

|  | Patient |  |  |  |  |  |  |  |  |  |
|---|---|---|---|---|---|---|---|---|---|---|
| Model | 3 | 11 | 12 | 15 | 37 | 58 | 77 | 87 | 91 | 100 |
| Statistical PINN (70/10/20) | 0.50 | **−0.03** | −8.21 | **0.88** | **0.83** | **0.90** | **0.82** | **0.51** | **0.60** | **0.66** |
| Retrained tM-GLV baseline | **0.67** | −0.18 | **0.15** | −1.14 | 0.71 | 0.67 | **0.83** | −21.04 | 0.12 | −0.19 |

## 4 Conclusion

The effectiveness of reinforcement learning for cancer treatment depends on how well its underlying simulator represents patient dynamics. This study evaluated physics-informed neural networks (PINNs) as a forecasting foundation for personalized intermittent androgen deprivation therapy, combining population learning with patient-specific representations and mechanistic constraints. By reserving the final 20% of each patient’s trajectory for testing, the study directly evaluated forecasting beyond the observed history used for training and validation. The results demonstrate promising forecasting performance relative to the chronologically retrained tM-GLV baseline, supporting further development of PINNs as patient-informed simulators for adaptive treatment.

Population training improved aggregate one-step prediction performance over the selected single-patient configuration, motivating the use of shared patterns in PSA dynamics to support individual predictions. Roll-forward training extended this approach to forecasting without repeatedly supplying observed future PSA. In the ten-patient comparison, the statistically sampled PINN configuration achieved a held-out *R*^2^ above 0.8 for 4 patients, compared with 1 patient for the tM-GLV model. Performance varied across patients and sampling strategies, and fine-tuning did not consistently improve on the pretrained population model. The favourable forecasts in several patients show the potential of a flexible neural model to capture trajectories that the selected mechanistic formulation represents less effectively.

The hybrid framework connects this predictive flexibility with a biological description of tumor growth and treatment response. Mechanistic constraints encode assumptions about the growth and competition of sensitive and resistant cell populations, while patient representations allow the network to learn differences in observed PSA and treatment histories. Together, these components provide a way to combine shared population patterns with individualized dynamics. Further work will examine how physics constraints and patient representations contribute to forecasting accuracy and simulated tumor dynamics.

This approach is relevant to adaptive therapy because treatment decisions alter both tumor burden and the competitive environment in which resistance develops. A patient-informed simulator could allow an RL agent to explore how treatment continuation, withdrawal, and resumption affect future trajectories, with the aim of controlling tumor growth while preserving sensitive cells that may suppress resistant expansion. Building on the simulator-based treatment optimization investigated by Gallagher et al. and Lu et al. [5, 6], the present work strengthens the forecasting component on which such policy exploration depends. Learned patient representations offer a route for conditioning simulated outcomes on individual histories while drawing on information from the wider cohort. The next step is to evaluate responses under alternative treatment schedules and use the refined simulator to train and assess personalized policies.

Overall, these findings support PINNs as a promising bridge between clinical time-series forecasting and mechanistic simulation for adaptive cancer therapy. The chronological evaluation establishes a foundation for predicting later PSA dynamics from an observed patient history. Extending this framework to adaptation from an initial treatment cycle or a small number of early visits would move it closer to prospective use, where treatment decisions must be made with limited information. This offers a path toward scalable, individualized simulation environments for investigating treatment schedules that seek to delay progression and reduce treatment burden.

## Footnotes

1 Details of the roll-forward model are in Section 2.7

2 Depot here denotes the constant, baseline release rate maintained by the tissue reservoir following the initial transient flare and stabilization phases.

† Not used in one-step PINNs, only in roll-forward models. Details of roll-forward models are in section 2.8

## Notes

### Competing Interest Statement

The authors have declared no competing interest.

## References

[1] Bruchovsky N, Klotz L, Crook J, Malone S, Ludgate C, Morris WJ, et al. Final results of the Canadian prospective phase II trial of intermittent androgen suppression for men in biochemical recurrence after radiotherapy for locally advanced prostate cancer: clinical parameters. Cancer. 2006;107(2):389–395. 10.1002/cncr.21989

[2] Zhang J, Cunningham J, Brown J, Gatenby R. Evolution-based mathematical models significantly prolong response to abiraterone in metastatic castrate-resistant prostate cancer and identify strategies to further improve outcomes. eLife. 2022;11:e76284. 10.7554/eLife.76284

[3] Strobl MAR, West J, Viossat Y, Damaghi M, Robertson-Tessi M, Brown JS, et al. Turnover modulates the need for a cost of resistance in adaptive therapy. Cancer Research. 2021;81(4):1135–1147. 10.1158/0008-5472.CAN-20-0806

[4] Engelhardt D. Dynamic control of stochastic evolution: a deep reinforcement learning approach to adaptively targeting emergent drug resistance. Journal of Machine Learning Research. 2020;21(203):1–30. https://jmlr.org/papers/v21/19-546.html

[5] Gallagher K, Strobl MAR, Park DS, Spoendlin FC, Gatenby RA, Maini PK, Anderson ARA. Mathematical model-driven deep learning enables personalized adaptive therapy. Cancer Research. 2024;84(11):1929–1941. 10.1158/0008-5472.CAN-23-2040

[6] Lu Y, Chu Q, Li Z, Wang M, Gatenby R, Zhang Q. Deep reinforcement learning identifies personalized intermittent androgen deprivation therapy for prostate cancer. Briefings in Bioinformatics. 2024;25(2):bbae071. 10.1093/bib/bbae071

[7] Tseng HH, Luo Y, Cui S, Chien JT, Ten Haken RK, El Naqa I. Deep reinforcement learning for automated radiation adaptation in lung cancer. Medical Physics. 2017;44(12):6690–6705. 10.1002/mp.12625

[8] Zhang J, Cunningham JJ, Brown JS, Gatenby RA. Integrating evolutionary dynamics into treatment of metastatic castrate-resistant prostate cancer. Nature Communications. 2017;8:1816. 10.1038/s41467-017-01968-5

[9] Sung H, Ferlay J, Siegel RL, Laversanne M, Soerjomataram I, Jemal A, Bray F. Global cancer statistics 2020: GLOBOCAN estimates of incidence and mortality worldwide for 36 cancers in 185 countries. CA: A Cancer Journal for Clinicians. 2021;71(3):209–249.

[10] Crawford ED, Heidenreich A, Lawrentschuk N, et al. Androgen-targeted therapy in men with prostate cancer: evolving practice and future considerations. Prostate Cancer and Prostatic Diseases. 2019;22(1):24–38. 10.1038/s41391-018-0079-0

[11] Holohan C, Van Schaeybroeck S, Longley DB, Johnston PG. Cancer drug resistance: an evolving paradigm. Nature Reviews Cancer. 2013;13(10):714–726. 10.1038/nrc3599

[12] Gatenby RA, Silva AS, Gillies RJ, Frieden BR. Adaptive therapy. Cancer Research. 2009;69(11):4894–4903.

[13] Lieberman R. Evidence-based medical perspectives: the evolving role of PSA for early detection, monitoring of treatment response, and as a surrogate end point of efficacy for interventions in men with different clinical risk states for the prevention and progression of prostate cancer. American Journal of Therapeutics. 2004;11(6):501–506.

[14] Raissi M, Perdikaris P, Karniadakis GE. Physics-informed neural networks: a deep learning framework for solving forward and inverse problems involving nonlinear partial differential equations. Journal of Computational Physics. 2019;378:686–707. 10.1016/j.jcp.2018.10.045

[15] Cho K, van Merriënboer B, Gulcehre C, Bahdanau D, Bougares F, Schwenk H, Bengio Y. Learning phrase representations using RNN encoder–decoder for statistical machine translation. In: Proceedings of the 2014 Conference on Empirical Methods in Natural Language Processing (EMNLP). 2014:1724–1734. 10.3115/v1/D14-1179

[16] Balk SP, Ko YJ, Bubley GJ. Biology of prostate-specific antigen. Journal of Clinical Oncology. 2003;21(2):383–391. 10.1200/JCO.2003.02.083

[17] Shimada T, Aihara K. A nonlinear model with competition between prostate tumor cells and its application to intermittent androgen suppression therapy of prostate cancer. Math Biosci. 2008 Jul-Aug;214(1-2):134–9. doi: 10.1016/j.mbs.2008.03.001. Epub 2008 Mar 13. PMID: 18420226.

[18] Gunnarsson EB, D. S, Leder K, Foo J. Understanding the role of phenotypic switching in cancer drug resistance. J Theor Biol. 2020;490:110162. doi: 10.1016/j.jtbi.2020.110162.

